# A framework for annotation of antigen specificities in high-throughput T-cell repertoire sequencing studies

**DOI:** 10.1101/676239

**Authors:** Mikhail V Pogorelyy, Mikhail Shugay

## Abstract

Recently developed molecular methods allow large-scale profiling of T-cell receptor (TCR) sequences that encode for antigen specificity and immunological memory of these cells. However, it is well known, that the even unperturbed TCR repertoire structure is extremely complex due to the high diversity of TCR rearrangements and multiple biases imprinted by VDJ rearrangement process. The latter gives rise to the phenomenon of “public” TCR clonotypes that can be shared across multiple individuals and non-trivial structure of the TCR similarity network. Here we outline a framework for TCR sequencing data analysis that can control for these biases in order to infer TCRs that are involved in response to antigens of interest. Using an example dataset of donors with known HLA haplotype and CMV status we demonstrate that by applying HLA restriction rules and matching against a database of TCRs with known antigen specificity it is possible to robustly detect motifs of an epitope-specific responses in individual repertoires. We also highlight potential shortcomings of TCR clustering methods and demonstrate that highly expanded TCRs should be individually assessed to get the full picture of antigen-specific response.

## Introduction

Immune repertoire profiling technology (AIRR-Seq, [1]) is an efficient technique that can be employed to study the structure and dynamics of the adaptive immune system. AIRR-Seq makes it possible to characterize the structure of both naive and antigen-experienced T-cell receptor (TCR) repertoires [2]–[4], tumor infiltrating T-cells [5], TCRs related to autoimmunity [6], leading to numerous downstream applications in both basic and applied immunological research [7]. While novel single-cell RNA sequencing methods allow coupling individual T-cell clones to their phenotype and function using their gene expression profiles [8], the actual antigen specificity (i.e. the set of antigens that can be potentially recognized by a given TCR) remains a mystery for most of the T-cells observed by high-throughput profiling. Even with deep repertoire profiling, the number of unique TCR variants obtained from MHC-multimer positive T-cell fraction is usually below 10^4^ [9], dwarfed by the highly conservative estimate of 10^8^ for the diversity of TCR beta chain [10].

Recent developments in the field of bioinformatic analysis of AIRR-Seq data are aimed at providing a mean for annotation of TCR repertoires with predicted antigen specificities. For example, McPAS-TCR database [11] lists pathogen- and disease-associated TCRs and VDJdb database [12] features a large set of TCRs with experimentally verified epitope specificities and their MHC restrictions. Existing computational methods for TCR repertoire annotation allow both matching against a database of known antigen specificities [12], [13] and clustering of TCR sequences for *de novo* motif detection [4], [14]. Annotation of a large number of TCR repertoires from healthy donors [15], [16] demonstrates both high variance of frequencies of epitope-specific T-cells and the imprint of past and ongoing pathogen encounters. Thus, *de novo* discovery of T-cells associated with antigens of interest or certain disease appears to be a hard problem, complicated by the biases in the structure of the naive (unperturbed) TCR repertoire [17], presence of existing clonal expansions specific to unrelated pathogens and high number of false positives that result from the extremely high diversity of the TCR repertoire.

In the present paper, we describe a general framework that can be used to infer sets of T-cells specific to antigens of interest using AIRR-Seq data and TCR neighborhood enrichment algorithms (ALICE and TCRNET). We discuss how various biases of AIRR-Seq datasets can be handled using proper experimental design and give a theoretical basis for the proper application of methods that are based on the probabilistic model of VDJ rearrangement. Using an example dataset of individual human TCR repertoires we demonstrate the capability of the framework to infer HLA-restricted antigen-specific responses, discuss possible modifications of the proposed method and expose potential shortcomings of the existing methodology that should be taken into account when running antigen-specific TCR inference.

## Materials and Methods

### AIRR-Seq data analysis

Six samples were selected from a large TCRbeta repertoire sequencing dataset published by Emerson *et al.* [18]. The samples were chosen based on HLA matching, they include a CMV- and CMV+ donors and 4 controls with CMV- status (see **Figure 3** for sample IDs). Short nucleotide sequences covering CDR3 region together with Variable (V) and Joining J) gene parts were then re-aligned with MiXCR software [19] to produce clonotype tables compatible with VDJtools software [20] and to resolve cases with missing V/J allele calls.

**Figure 1.**
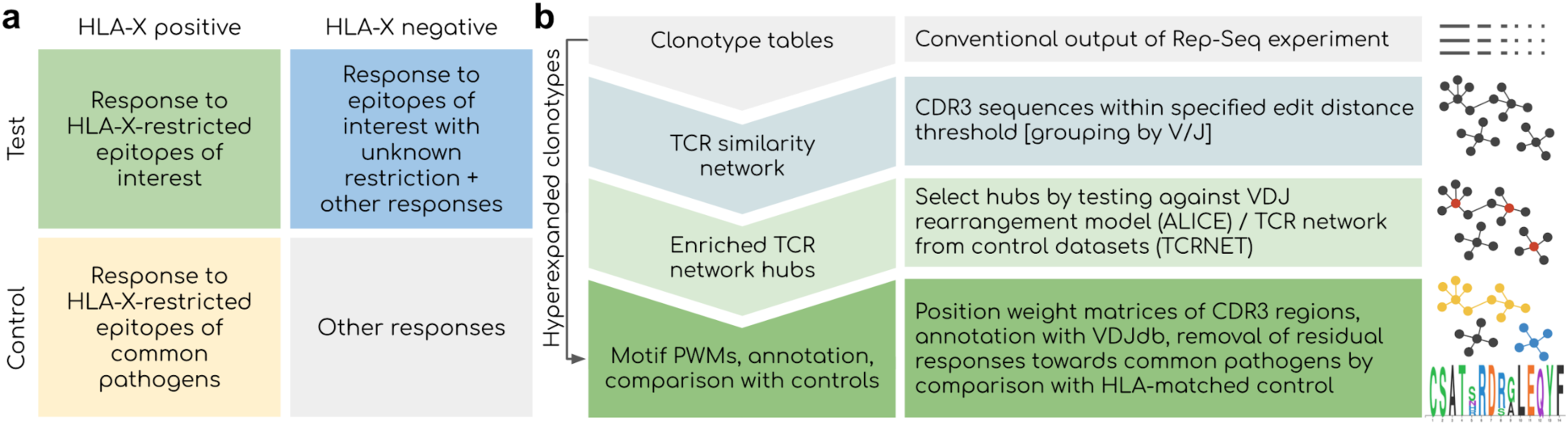
Overview of the proposed framework. **a.** Considerations for experimental design. Controlling for HLA type is critical as the response towards epitopes linked to the disease of interest is shaped by HLA binding restrictions: donors with distinct HLA can have a completely different response linked to the same disease. Using a control set with matched HLAs is also critical as a response to present and past common infections will be present among all donors. **b.** Outline of the bioinformatic analysis pipeline. Network-based analysis is aimed at detecting groups of T-cell clonotypes with homologous TCRs that were expanded during antigen-specific response. Statistical testing offered by ALICE/TCRNET methods is required to control for the intrinsic structure of TCR repertoire shaped by the process of VDJ rearrangement and presence of common pathogens (TCRNET case). Downstream annotation with VDJdb can be used to infer actual antigen specificities of TCR clusters and to remove common antigen-specific responses in case of a study aimed at detecting a response to novel/rare pathogens and self-antigens. Hyperexpanded clonotype should be considered even in case they are not included in homologous TCR clusters for the reasons described later in the following sections of the paper.

**Figure 2.**
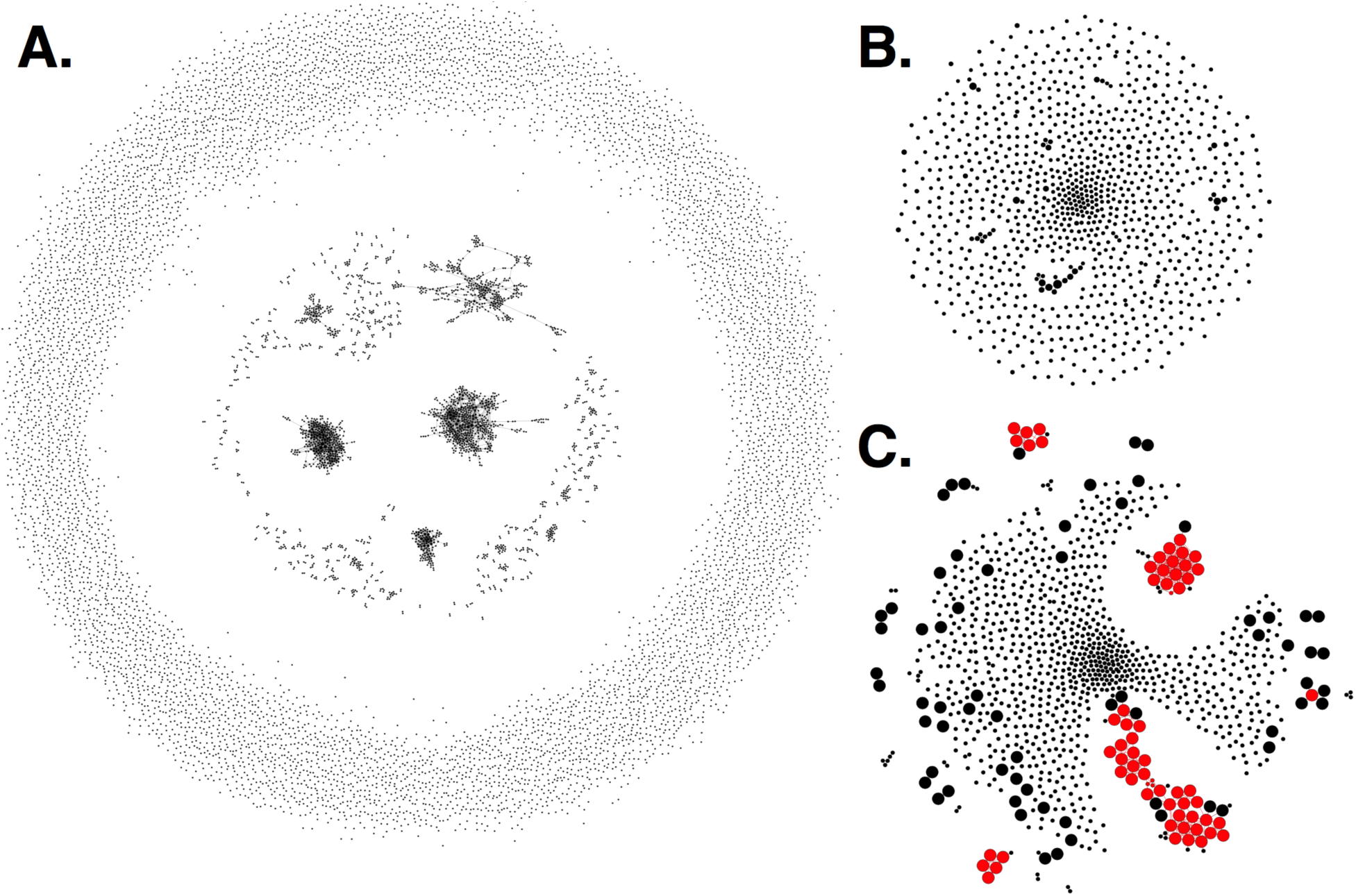
Rep-Seq sample simulated according to the VDJ rearrangement model. **a.** The similarity network of 10,000 randomly generated TRBV7-6/TRBJ1-4 TCRs. Each vertex shows an individual clonotype, edges indicate Hamming distance of 1 or less between CDR3 amino acid sequences. **b.** A TCR similarity network of 1,000 clonotypes randomly sampled from (**a.**) modeling uniform selection from the repertoire. ALICE algorithm identifies no hits (clusters) in this network. **c.** Modeling an antigen-driven selection by a 100-fold increase in the sampling probability of 50 randomly selected clonotypes and all their observed neighbours from (**a.**). ALICE hits are shown in red. Vertex size shows antigen-driven expansion fold in the initial repertoire. ALICE algorithm identified as significant hits 60 clonotypes (true positives, large red circles) out of 126 expanded in this simulation and 4 unexpanded clonotypes belonging to enriched clusters (false positives, small red circles), the rest 870 clonotypes are thus true negatives. While there are some clusters of cooperatively expanded similar clonotypes, there are also a lot of expanded singletons, which have no neighbours (large black circles, false negatives).

**Figure 3.**
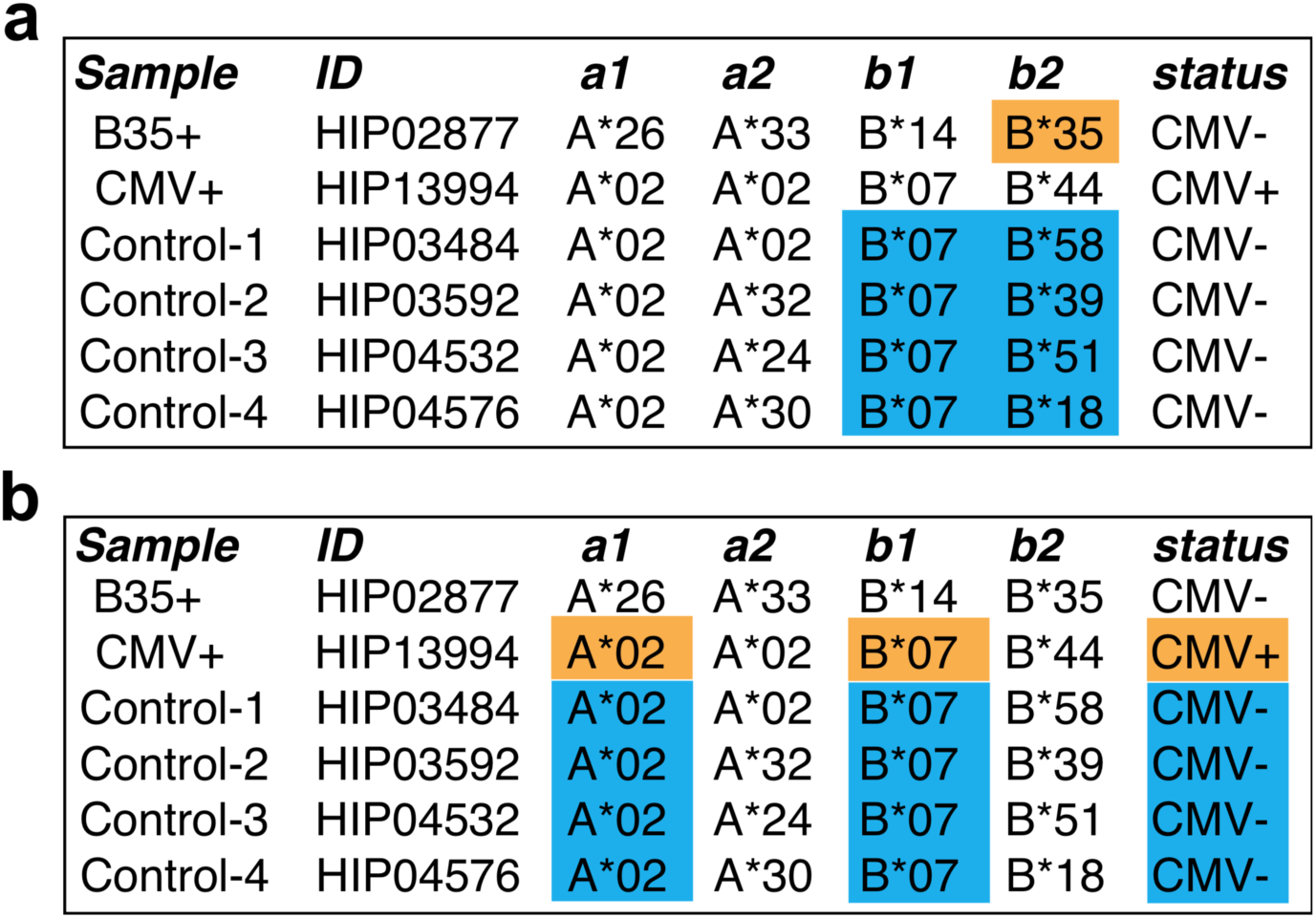
The design of a benchmark experiment. **a.** A case study of responses restricted to HLA-B*35 allele. B*35-positive donor is highlighted with orange, control samples are selected in the way that there is no B*35 allele are highlighted with blue. Note that the B*35-positive donor is CMV-negative. **b.** A case of CMV-specific response linked to HLA-A*02 and HLA-B*07 allele. A CMV+ donor (orange) is compared to CMV-controls (blue), with A*02 and B*07 alleles matched across all donors. In this case, one expects to find specific TCRs recognizing CMV epitopes presented by A*02 and B*07. Selected samples from the Emerson *et al.* [18] study were used.

### VDJ rearrangement simulation, network analysis and repertoire annotation

Random TCRbeta sequences were simulated using OLGA software [21] with default VDJ rearrangement model parameters and V/J allele sequences. TCR similarity networks were constructed by allowing a single substitution (a Hamming distance of 1) in CDR3 amino acid sequences. Neighborhood size (degree) enrichment of TCR similarity network nodes was tested against VDJ rearrangement model using ALICE algorithm [4]. Minimal number of neighbours was set to 2, Q selection factor was set to 1(no thymic selection) for the analysis of sequences generated with OLGA (see Fig 2), and to 9.41 (default) for the analysis of HSCT dataset (see Fig. 5).. Node neighborhood enrichment test against a pooled control dataset of real TCR repertoires was performed using the TCRNET algorithm implemented in the VDJtools software [2], [20]. TCR repertoire annotation was performed using the VDJdb database [12] with a single substitution allowed in the CDR3 amino acid sequence using VDJmatch software (https://github.com/antigenomics/vdjmatch).

**Figure 4.**
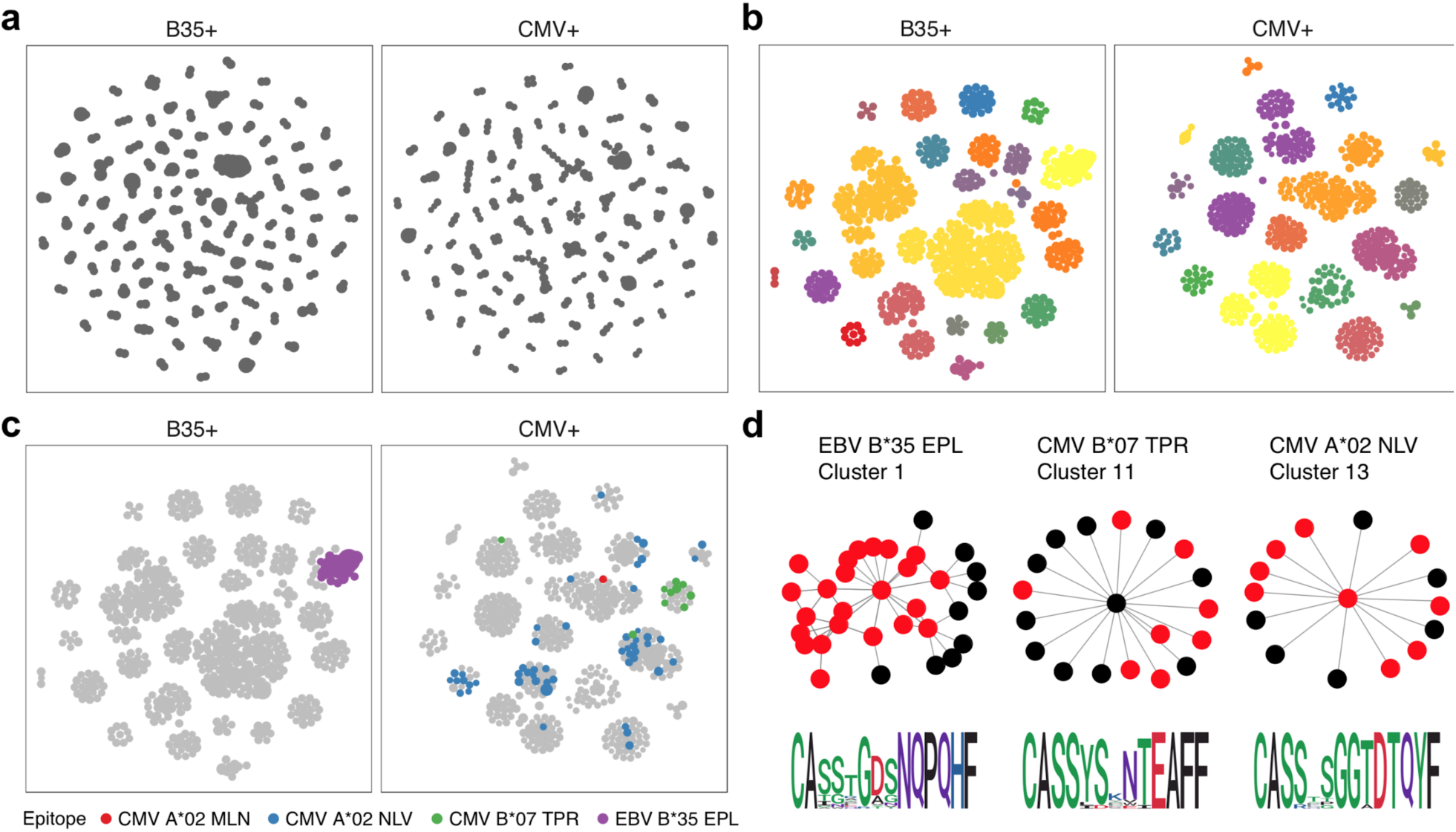
Results of proposed framework when applied to the benchmark experiment. **a.** A TCR similarity network built for top 3,000 TCRs present in B35+ and CMV+ samples. **b.** Selected network hubs for the TCR similarity network built on top of TCRs enriched in neighbour sequences. Each connected component is colored with a distinct color. c. VDJdb annotations of network hubs based on donor status and HLA restriction. Color shows TCR epitope specificity. **d.** TCR motifs inferred for selected TCRNET clusters. Graph shows the structure of the connected component of a given cluster, red nodes represent TCRs annotated with a given epitope specificity according to VDJdb (1 amino acid substitution allowed for CDR3aa). Sequence logos of CDR3aa sequences are given at the bottom of each cluster. TCR similarity networks are created using a Hamming distance threshold of 1 for CDR3aa sequences. Multidimensional scaling (MDS)-based graph layout is used for graph visualization in two dimensions.

**Figure 5.**
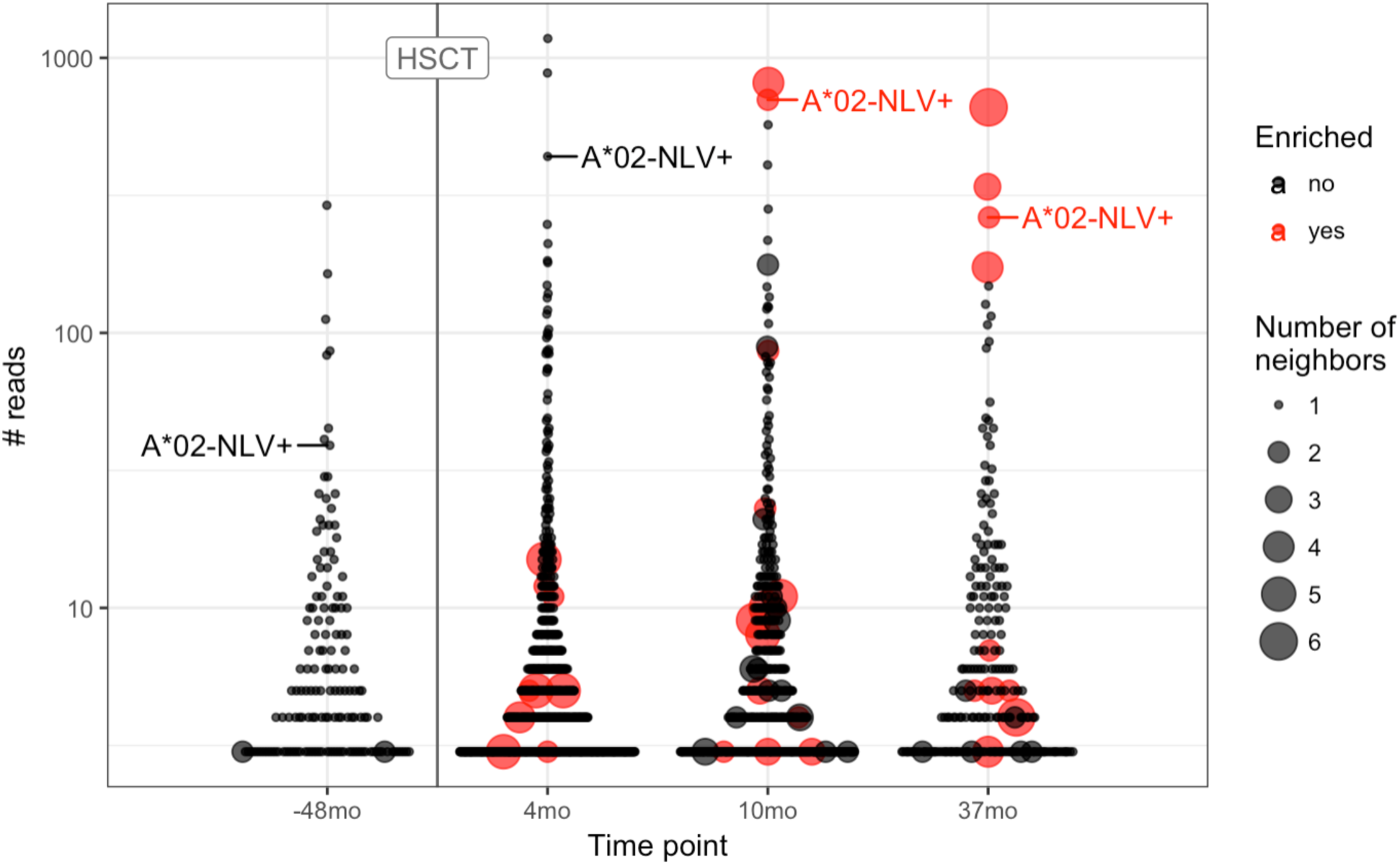
TCR motifs inferred by ALICE and tracking of CMV-specific TCR expansion for the hematopoietic stem cell transplant (HSCT) experiment. Beeswarm plots show the distribution of the number of reads associated with TCR clonotypes prior and after HSCT. The A*02-NLV-specific TCR variant with ‘CASSLAPGATNEKLFF’ CDR3aa sequence is highlighted for both the sample prior to HSCT (−38mo) and later samples (4, 10 and 37mo). Red dots show the clonotypes that have unexpectedly high number of neighbors as inferred by ALICE (P < 0.001 after Benjamini-Hochberg multiple testing correction), point size reflects the number of neighbor sequences. Only TCRs supported by at least 3 sequencing reads are shown.

### Code and data availability

Example datasets and an R markdown notebook for the described framework can be accessed at (https://github.com/antigenomics/scair-tutorial-2018). Running the code requires installing several R packages listed in the notebook and Java 1.8+. The code was tested and runs without problems on a Unix system with a 4 core Intel CPU and 6GB RAM.

## Results

### Considerations for experimental design and the analysis pipeline

There are several factors that should be controlled for when searching for TCR motifs associated with a certain treatment or disease (**Figure 1a**). Firstly, HLA restriction is the major factor that shapes the entire response: TCR motifs result from a response targeting certain epitope and are very likely to be absent in case some of the donors do not have a specific HLA haplotype even when there are no other differences between donor phenotypes. Thus, HLA typing is a prerequisite for any AIRR-seq study that aims at detecting TCR motifs and both test and control cohorts should be carefully balanced according to HLA frequency.

Another factor to account for is the imprint of past infections that is subject to HLA restriction [15], [16]. Multiple clonal expansions related to common pathogens can be detected across a partially HLA-matched set containing patients and healthy controls that are unrelated to the studied case. Therefore, a large HLA-matched cohort of healthy donors is required to filter out TCR motifs associated with common infections such as EBV.

Finally, the features of the unperturbed TCR repertoire structure itself should be considered, as the repertoire is heavily shaped by the process of VDJ rearrangement. The fact that the VDJ rearrangement process can be described by a relatively simple probabilistic model makes it possible to accurately predict population frequencies of specific TCRs [15]. However, huge differences in epitope-specific TCR frequencies can lead to the detection of potentially irrelevant high-frequency (public) TCRs in case the cohort size is relatively small as illustrated in [22]. Moreover, those public TCRs are the basis for large hubs of TCR similarity network [17], therefore they can be mistaken for real homologous clonal expansions. The means to handle this factor are described in the next section and form the basis of the proposed TCR motif inference framework (**Figure 1b**).

### Theoretical basis for TCR sequence motif inference

There are three assumptions that make it possible to detect a set of homologous TCRs that are involved in an ongoing antigen-specific response from AIRR-seq data:

**1a)** TCR rearrangement process follows probabilistic model of Murugan *et al.* [23]. This assumption allows to compute the expected incidence rate of a given TCR, which is in a good agreement with observed results [15] in case a single amino acid substitution is allowed in the CDR3 sequence.

**1b)** As it follows from [23], average TCR sequence is extremely rare: median rearrangement probability across all TCRs is ∼10^−12^. So given one samples ∼10^6^ T-cells from a total of ∼10^11^ T-cells in human peripheral blood, the expected count for a TCR clonotype is <<1. This allows us to model TCR sampling using Poisson distribution and is required for point 3).

**2)** There are multiple T-cells with homologous TCRs that recognize the same epitope in an individual. As can be observed in MHC-multimer sequencing studies (see e.g. [24]), it is typical for the set of multimer+ cells to contain groups of homologous TCRs. Although, it is still possible that the response to a given epitope is either monoclonal or utilizes a set of distinct TCR sequences, in which case it is not possible to infer any motif.

**3)** Antigen-specific T-cells expand upon antigen exposure and rare variants go above the detection boundary 1b). This effect allows us to run a neighborhood enrichment test to detect the antigen-driven response without relying on clonotype frequency statistics and pre-exposure control samples.

The assumption 1) can be utilized to build a control TCR similarity network that recaptures biases of the VDJ rearrangement process. An example of a network of 10,000 randomly generated TCRs constructed by connecting CDR3aa regions that differ by a single substitution is given in **Figure 2a**. This network reveals a complex structure with multiple hubs that, as previously shown in [17], are enriched in ‘public’ TCR variants. It is necessary to note that disconnected hubs may arise, in part, due to the fact that TCRs with different CDR3 length are not connected when using Hamming distances. Allowing for indels or more substitutions, on the other hand, leads to larger hubs at the cost of greatly increasing the number of false-positives (see [12]).

These theoretic assumptions provide a basis for usage of TCR neighborhood enrichment tests implemented in ALICE and TCRNET algorithms [2],[4]. The difference between the latter two is that ALICE uses VDJ rearrangement model described by Murugan *et al.* as a control, while TCRNET utilizes pools of healthy control samples as background (negative) set of clonotypes. Thus, the former tests against VDJ rearrangement biases and thymic selection biases defined by the Q-factor [4], while the latter implicitly tests against thymic and peripheral selection biases and common pathogen-specific responses based on the structure of the control TCR set.

In a random sample of 1000 TCRs one can still observe some of those hubs, however, those are smaller and lack any TCRs enriched with neighbours according to the ALICE algorithm (**Figure 2b**). We next simulate antigen-specific clonal expansion by sampling 50 random TCRs and their observed one mismatch neighbours at 100x higher rate. This simulation leads to **Figure 2c** in which ALICE detects several enriched clusters, though there are multiple expanded TCRs having no neighbours (monoclonal expansions). Note that the latter are naturally occurring and were previously observed in multiple studies that involve tetramer-based enrichment of antigen-specific T-cells [13]; a detailed description of such cases is provided in the following sections.

### Inferring CMV-specific and B35-restricted responses

Straightforward annotation of selected samples (CMV+, B35+ and 4 pooled controls) by querying the VDJdb database with 1 substitution allowed (see **Materials and Methods**) results in a huge variety of antigen specificities (**Supplementary Figure 1**). The latter include HIV- and HCV-specific clonotypes that are not expected for systematically healthy individuals and it is hard to tell the overall difference observed in CMV and B35 samples with respect to control. Applying HLA (restricting to HLA-B*35 for B35 sample and HLA-A*02/HLA-B*07 for CMV sample) and pathogen restriction rules (CMV for CMV+ sample) that follow from our experimental design greatly reduces the complexity of observed results (**Supplementary Figure 2**). However, while the presence of EBV-related B*35:EPL clonal expansions is evident for the B*35 donor, it is hard to tell whether the CMV-specific clonal expansions are significantly different from control for the CMV sample.

We therefore ran the *de novo* motif discovery procedure for B35 and CMV samples using the TCRNET algorithm and specifying the background dataset to be the pool of 4 control samples (see **Materials and Methods**). Notably, there is a correlation of TCR neighbor enrichment rate with the overall expansion of those TCRs (**Supplementary Figure 3**). As one can see from **Figure 4ab** the TCR similarity network selected by TCRNET is substantially different from the one built using top (by frequency) 1000 clonotypes from those samples. Namely, while the latter resembles the unperturbed network of random VDJ rearrangements with power law distribution of hub sizes, the network observed in **Figure 4b** has a more uniform hub size distribution. By annotating the resulting network against VDJdb (**Figure 4c**) one can see the presence of network hubs that have a huge fraction of associated clonotypes annotated with the same antigen specificity (**Table 1**). We can, therefore, derive position weight matrices for specific responses (**Figure 4d**) and build corresponding TCR motifs. Notably, this way we do re-assign many clonotypes of unknown specificity to the predicted specificity of their hubs and can further extend our knowledgebase of TCRs of known specificity with these predictions.

**Table 1.**
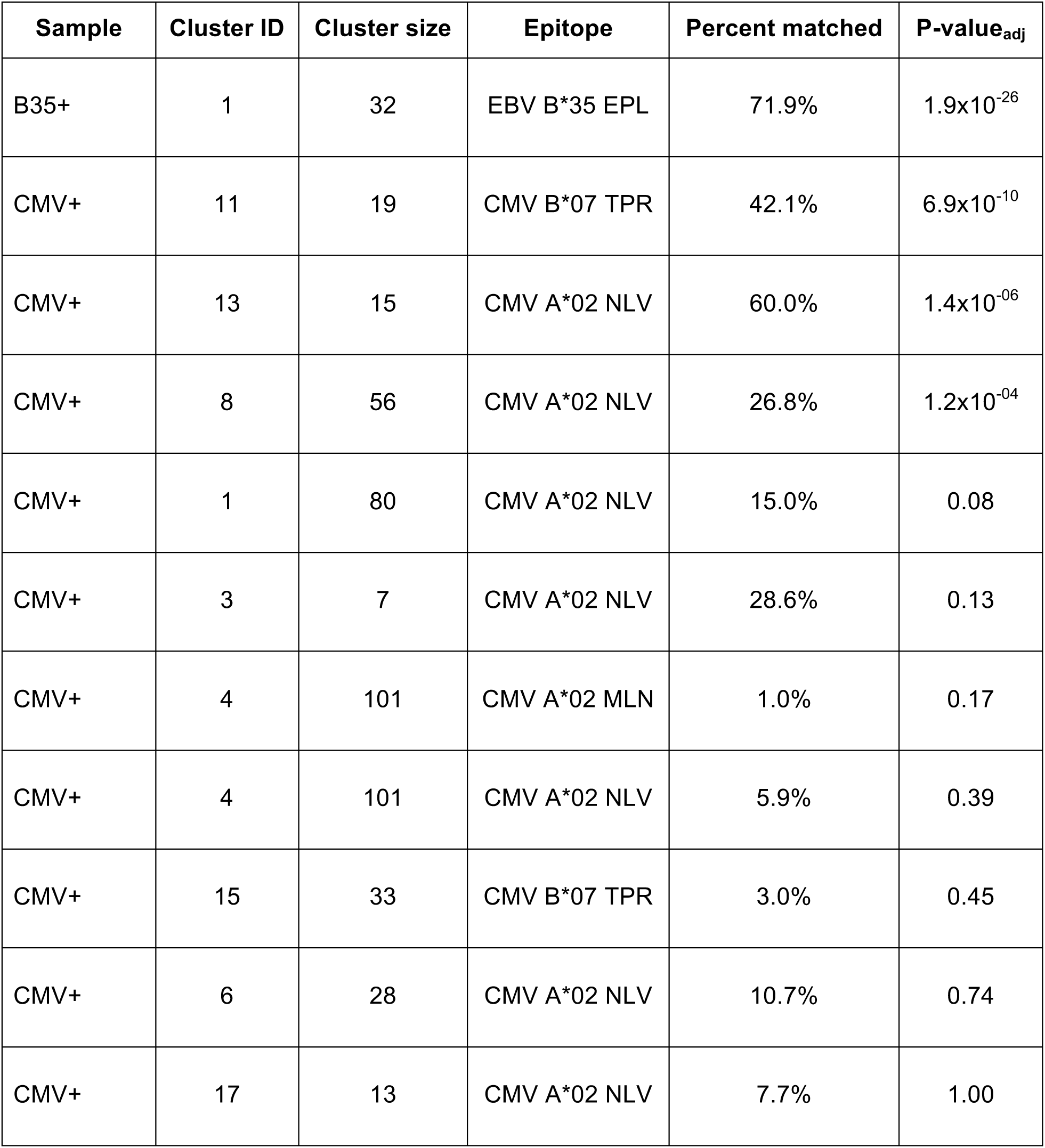
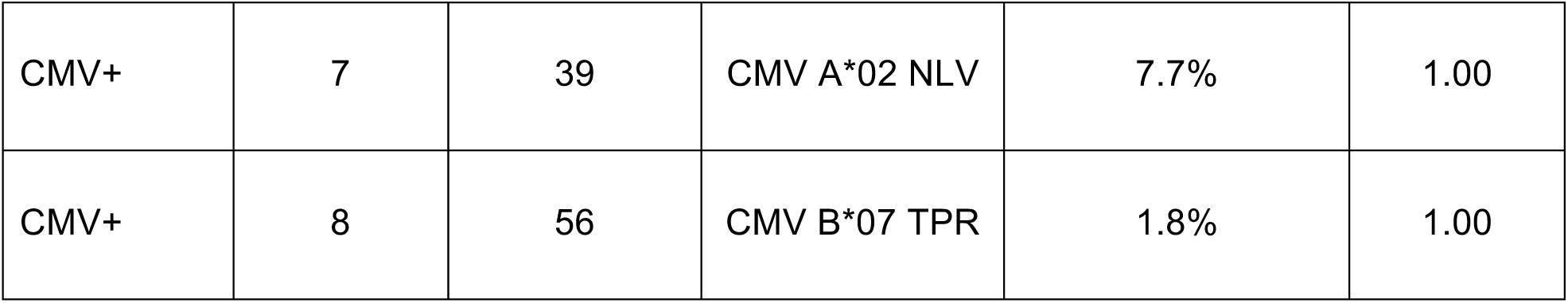
VDJdb annotation results for TCRNET clusters. The table shows the percent of matches to a given epitope in a given cluster according to VDJdb database (1 amino acid substitution allowed for CDR3aa). P-values for the frequency of matches were computed based on Binomial distribution using overall annotation rate across the sample, the number of specific TCR matches and the size of a given cluster, p-values were adjusted for multiple testing. Clusters that have no VDJdb matched are not shown.

### Exploring the case of dominant clonal expansions

While we were able to obtain a set of high-confidence TCR motifs in the previous section, a deeper exploration of the dataset, however, reveals that there is a huge fraction of T-cells that were missed by our analysis. Namely, when exploring VDJdb hits that are associated with large clonal expansions, one can see that there are certain large clonal expansions that are exactly matching to CMV specific clonotypes, yet do not fall into any of the listed motifs (**Supplementary Figure 1**).

To further highlight this issue we ran the pipeline for hematopoietic stem cell transplant (HSCT) time course data [25] that is known to result in large CMV-specific clonal expansions (**Figure 5**). We followed the fate of the A*02 NLV specific TCRbeta CDR3aa sequence CASSLAPGATNEKLFF to trace the corresponding response. Corresponding TCR clonotype is detected in all time points, both prior and after HSCT, where it reaches top 3 of expanded clonotypes. This clonotype is one of those that occupy homeostatic space following repertoire reset during HSCT and one would expect that homologous TCRs involved in CMV-specific response will follow it. However, while corresponding clonal expasion is evident both prior to HSCT and throughout the whole time course, homologous variants start to emerge and become detectable by the ALICE only after 10 months post-HSCT. This suggests that tracking of hyperexpanded clonotypes can be more sensitive than TCR neighborhood enrichment methods at early stages of response where the latter fail to detect a sufficient number of homologous TCRs.

## Discussion

As demonstrated using the example dataset the proposed framework can successfully detect both TCR motifs associated with specific pathogens and TCRs associated with a response towards epitopes restricted by a given HLA. Noting that EBV specific response detection is well expected due to the fact that most individuals are EBV-positive, it is necessary to mention that a multitude of unknown responses are detected for B35 sample and most CMV-specific TCR annotations are restricted to individual clusters in the CMV donor, suggesting that proposed method has both high precision in detecting known antigen-specific responses and high recall for detecting novel ones. As demonstrated above, the knowledge of donor HLAs can greatly simplify the analysis by narrowing down the list of potential specific TCR candidates. Currently, this seems the only way to combat the huge number of false positives that arise during VDJdb matching due to the immense diversity of TCR repertoires. On the other hand, the framework that we propose has the benefit of eliminating spurious matches that arise due to the presence of “public” clonotypes that can be shared across a wide range of samples by chance.

The proposed approach can be further extended to a wide range of applications beyond previously reported detection of antigen-specific response in yellow fever virus vaccination studies [26]. A successful application of a simpler VDJ rearrangement model-based approach that does not utilize TCR similarity networks to autoimmunity studies with strong HLA-linkage such as ankylosing spondylitis [6] suggests that our approach can be utilized for detecting autoimmunity-related TCR motifs. One can also apply the proposed approach to cancer studies, in case an overexpression of certain oncogenes or oncogenic isoforms is expected. For the latter, one should expect the presence of an immunogenic neoantigen that is both restricted to certain HLA allele and is characterized by overall low expression in healthy tissue.

We also suggest that the proposed framework can be used to expand the set of known antigen-specific TCRs that is currently negligibly small (∼10^4^ variants, [12]) compared to the overall repertoire diversity. Indeed, one can assign neighbours of enriched specific TCR clonotypes to the set of epitope-specific T-cells once corresponding homologous TCR clusters are detected in multiple donors. As some of the clusters have a statistically significant number of annotated TCRs even in the presence of a huge fraction of unannotated ones, one can expect to greatly increase the number of clonotypes in a database of TCRs with known specificity using those putative TCR variants.

One of the drawbacks of the proposed methodology follows from the fact, that in case when the HLA-disease association is relatively vague with no predominant HLA disease susceptibility (e.g. multiple sclerosis [27] or type 1 diabetes [28]) a huge cohort of donors featuring various HLAs should be used to detect potential TCR motifs for self-antigens. Another potential issue results from the presence of monoclonal expansions that are hard to cluster into inferred TCR motifs. The solution here is to treat all hyperexpanded clonotypes separately and rely on a database of TCRs with known antigen specificities to annotate those clonotypes. The proposed framework is designed for single-chain data, however, it is rather straightforward to extend it to paired alpha-beta TCR analysis: in case certain pairing information is available via, say, scRNA-seq, one can try to pair individual alpha and beta TCR motifs based on their co-occurrence in single-cell data.

Both the methodology and the interactive analysis notebook provided as a supplementary to this manuscript are easy to extend and adapt for post-analysis of various AIRR-seq datasets, we hope that the described framework will be of high utility for future exploratory AIRR-seq studies that aim at discovering novel antigen- and disease-specific TCR variants.

## Acknowledgements

Some parts of this work were originally presented during a tutorial session at the 2nd Meeting on Stochasticity and Control in Adaptive Immune Repertoires (Paris, 2018), the authors would like to thank the participants of this tutorial for their valuable feedback. This study was supported by Russian Science Foundation Grant No. 17-15-01495.

## Supplementary Figures and Tables

**Supplementary Figure 1.**
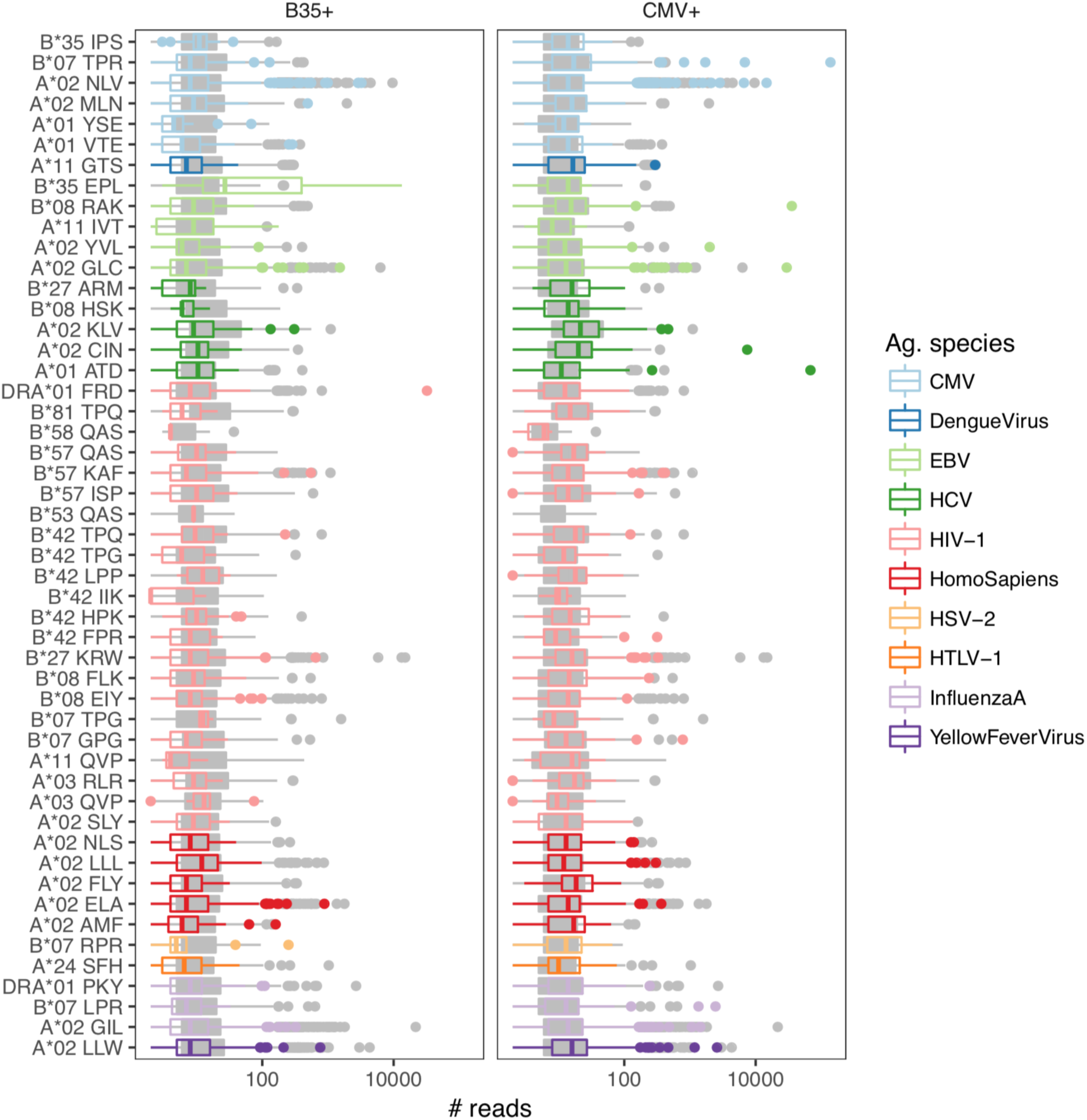
VDJdb annotation results for B35+ and CMV+ samples compared to pooled control dataset. Box plots of the number of reads associated with annotated TCRs for various epitope specificities. Epitope specificities are encoded as the restricting HLA (e.g. B*35) and first three amino acids of the epitope sequence (e.g. IPS). Boxplots are colored according to the parent species of the epitope. Grey boxplots show the frequency in pooled control samples. VDJdb annotation is performed with 1 amino acid substitution allowed for CDR3aa.

**Supplementary Figure 2.**
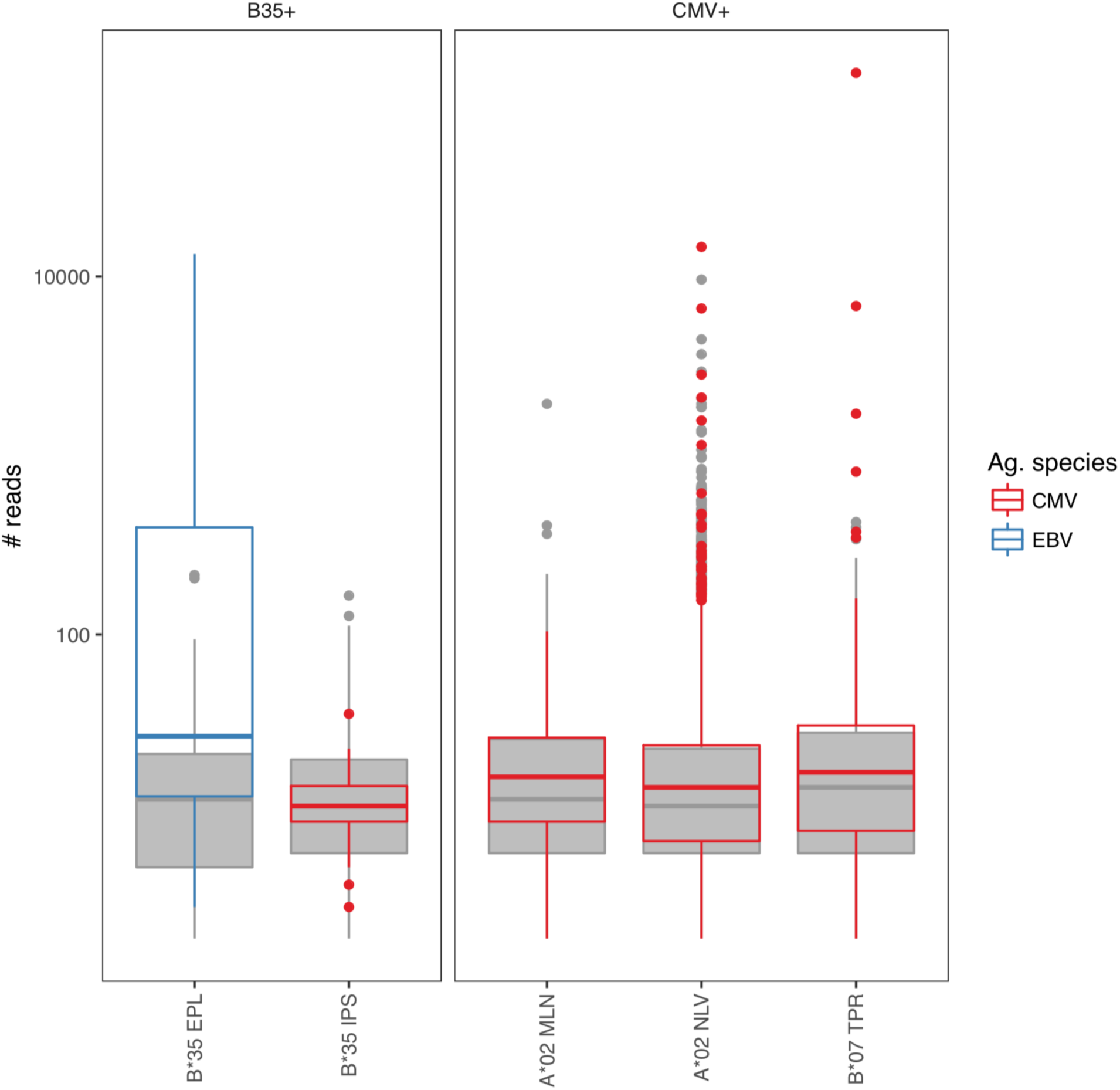
Epitope-specific TCR frequencies in B35+, CMV+ and pooled control samples for selected epitopes. Boxplots show TCR frequency in the sample. Epitopes were filtered according to donor status and HLA restriction: HLA-B*35 for B35+ sample and CMV+/HLA-B*07 or HLA-A*02 for CMV+ sample. Boxplots are colored by parent species of the epitope, grey boxplots show the TCR frequency in pooled control samples.

**Supplementary Figure 3.**
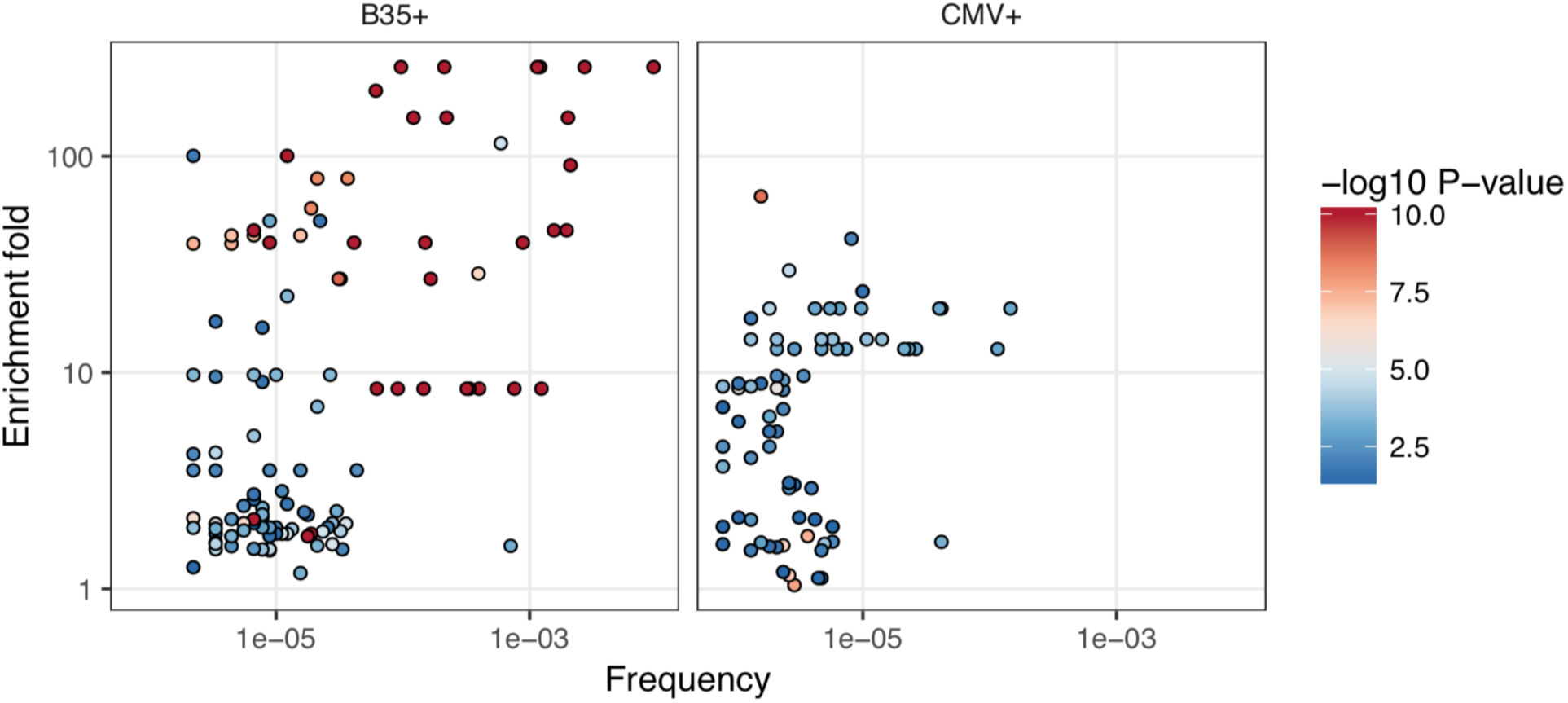
Comparing enrichment fold to TCR expansion rate. Scatter-plot comparing fraction of observed and expected neighbor sequences (enrichment fold, computed using TCRNET algorithm) and the T-cell expansion (TCR frequency in sample). Each point represents a TCR clonotype, points are covered by the enrichment fold P-value (TCRNET test). Significant correlation is observed for both B35+ (Spearman R=0.42, P=3×10^−6^) and CMV+ (Spearman R=0.31, P=8×10^−3^) samples.

**Supplementary Table 1.**
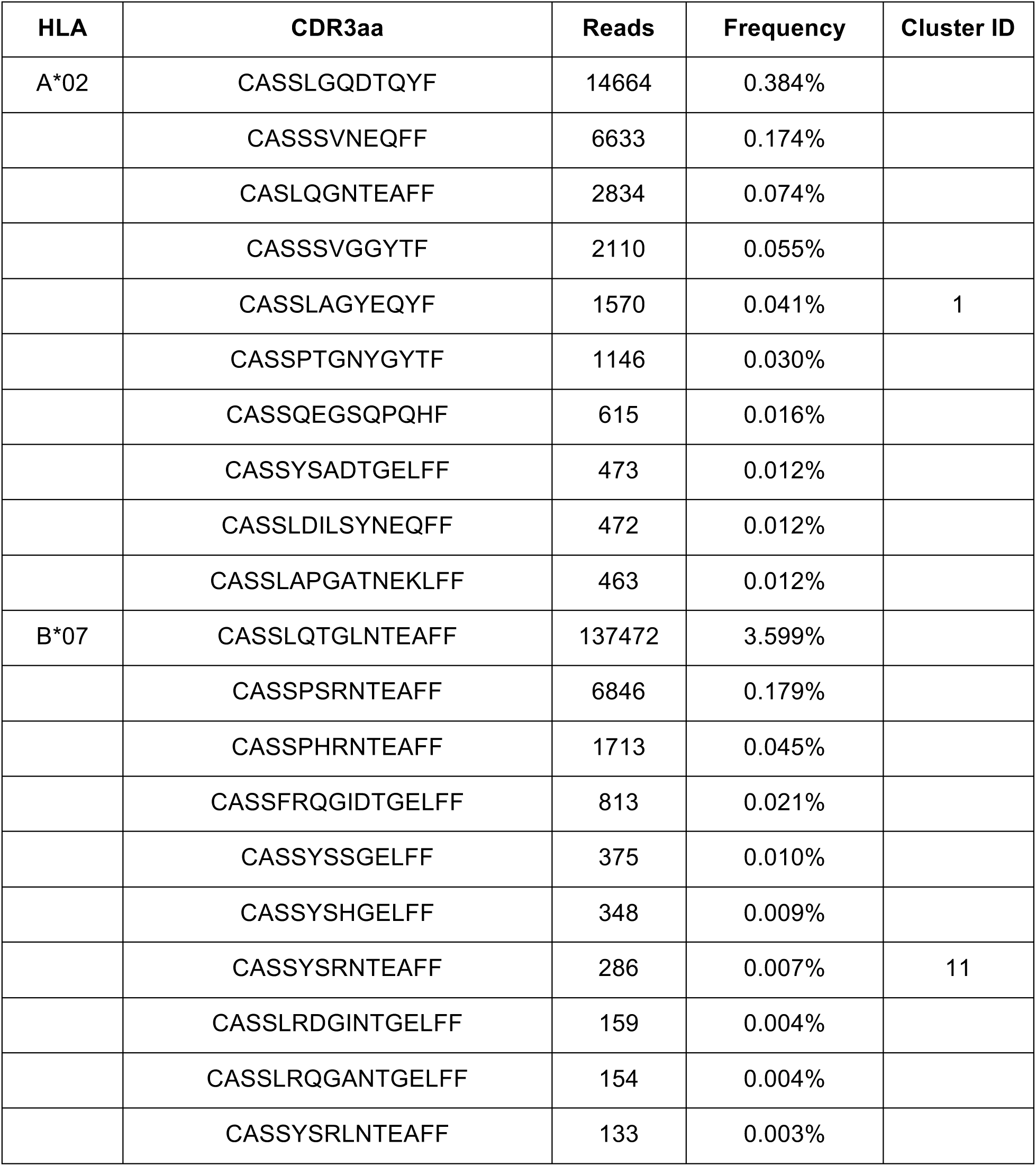
TCRNET clusters across top 10 most frequent specific TCRs annotated according to VDJdb. CMV-specific clonotypes with A*02 and B*07 HLA restriction are shown with corresponding number of reads. In case a TCR clonotype belongs to an inferred TCR cluster a cluster ID is provided.

## Notes

https://github.com/antigenomics/scair-tutorial-2018

